# Structural basis for assembly of human cN1A filaments

**DOI:** 10.64898/2026.11.20.768069

**Authors:** Fleur N. Brinkman, S. John Calise, Amrah Weijn, Josephine Molfino, Joel D. Quispe, Anke Rietveld, Megan K. Herbert, Annemarie van der Heijden, Evan Spruijt, Justin M. Kollman, Ger J.M. Pruijn

**Affiliations:** Department of Biomolecular Chemistry, Institute for Molecules and Materials, Radboud University, Nijmegen, The Netherlands; Department of Biochemistry, University of Washington, Seattle, WA, USA; Department of Neurology, Center for Neuroscience, Donders Institute for Brain, Cognition and Behaviour, Radboud University Medical Center, Nijmegen, The Netherlands; Department of Physical Organic Chemistry, Institute for Molecules and Materials, Radboud University, Nijmegen, the Netherlands

**Keywords:** Cytosolic 5’-nucleotidase 1A, Inclusion body myositis, Cytoophidia, Rods and rings, Filaments, Nucleotide metabolism, Protein-protein interactions, Cryo-EM

## Abstract

Human cytosolic 5’-nucleotidase 1A (cN1A), which catalyzes the hydrolysis of AMP to adenosine, is a prominent autoantigen in the progressive myopathy inclusion body myositis (IBM). Detection of anti-cN1A autoantibodies is a supportive criterion for IBM diagnosis, and mice immunized actively or passively with cN1A develop an IBM phenotype. However, how cN1A breaks immunotolerance is not yet known. Despite its clinical significance, the subcellular localization of cN1A is not well understood, and its structure has not been determined. Here, we show that human cN1A is most highly expressed in type 2 myofibers and tends to localize near the periphery of muscle fibers, but is either weakly expressed or undetectable in commonly used human cell lines. Ectopic expression of cN1A in such cells resulted in spontaneous assembly of cN1A into filamentous structures strikingly similar to metabolic filaments formed by the nucleotide biosynthesis enzymes IMP dehydrogenase (IMPDH) and CTP synthase (CTPS). However, cN1A filaments did not co-localize with IMPDH or CTPS filaments and were not responsive to compounds that induce IMPDH and CTPS assembly. Negative stain electron microscopy (EM) showed that purified recombinant cN1A protein spontaneously polymerizes, suggesting filament assembly in cells is driven by intrinsic properties of cN1A. We then determined the structure of the cN1A filament by cryo-EM, which revealed that cN1A monomers assemble end-on-end into a polar helical filament and allowed us to identify contacts mediating filament assembly. Structure-guided mutations designed to abolish filament assembly prevented cN1A filament formation in cells, establishing the structural basis for cN1A filament assembly.

## Introduction

Sporadic inclusion body myositis (IBM) is an acquired inflammatory and degenerative myopathy most commonly occurring in patients over 45 years of age. The disease is characterized by slowly progressive asymmetric weakness of proximal leg and distal arm muscles and dysphagia.^1^ Diagnosis is based on a set of defined clinical, serological, and histological criteria that are supported by newer criteria, including positive detection of autoantibodies against cytosolic 5’-nucleotidase 1A (cN1A; gene: NT5C1A),^2^ which were first reported in IBM patient sera in 2013.^3,4^ Anti-cN1A autoantibodies are present in 33-76% of patients with IBM, in other inflammatory myopathies like polymyositis (4-14%) and dermatomyositis (4-13%), and in autoimmune diseases such as systemic lupus erythematosus (6-21%) and Sjögren disease (7-19%).^5–8^ Anti-cN1A reactivity has been correlated with a more severe IBM phenotype and a higher mortality risk compared to anti-cN1A-negative IBM patients,^9^ although such correlations were not confirmed in other studies.^6,7,10^

Accumulation of cN1A has been detected in the muscles of IBM patients, in particular in the perinuclear region,^3,7,11,12^ surrounding rimmed vacuoles,^3,11,12^ near sarcoplasmic membranes, and in the sarcoplasm.^7^ In contrast to the relatively high cN1A expression levels in muscle tissue, cN1A is only poorly expressed in other tissues and human cell lines.^13^ Previously, while developing a novel immunofluorescence assay to detect anti-cN1A autoantibodies,^14^ we found that a GFP-tagged cN1A fusion protein spontaneously formed micron-scale filaments when expressed in the immortalized human HEp-2 cell line. This robust assembly of cN1A into large, visible filamentous structures enabled more sensitive and specific detection of anti-cN1A autoantibodies than existing enzyme-linked immunosorbent assays (ELISA).^14^

cN1A is an important enzyme in nucleotide metabolism, contributing to maintaining homeostasis by catalyzing the hydrolysis of AMP into adenosine, and potentially other nucleoside monophosphates, such as IMP, into their corresponding nucleosides.^13^ By light microscopy, cN1A filaments resemble filaments formed by other enzymes in nucleotide metabolism, including IMP dehydrogenase (IMPDH) and CTP synthase (CTPS), which catalyze the rate-limiting steps in *de novo* guanosine and cytidine nucleotide synthesis, respectively. The assembly of IMPDH and CTPS into micron-scale filamentous structures (also called “rods and rings” or “cytoophidia”) is well documented.^15,16^ Filament assembly in cells is a dynamic response to nucleotide deficiency and proliferative metabolic signaling,^17–20^ consistent with their *in vitro* biochemical function of resisting feedback inhibition to increase overall activity.^21–23^ In cell culture, IMPDH and CTPS filament assembly can be reversibly induced by various methods that perturb nucleotide metabolism.^24,25^ Interestingly, like cN1A, autoantibodies targeting IMPDH filaments (anti-rods/rings) were discovered over 15 years ago in patients with chronic hepatitis C virus infection treated with interferon and ribavirin.^26^

Here, we observed accumulation of cN1A at the periphery of myofibers in both IBM patients and healthy individuals. To further investigate cN1A assembly, we expressed cN1A in different human cell lines and found that it spontaneously forms filaments regardless of cell type, hinting that these filaments are likely related to cN1A accumulation in muscle tissue. We then purified recombinant human cN1A and demonstrated with negative stain electron microscopy (EM) that it has an intrinsic tendency to polymerize. Moreover, we determined a cryo-EM structure of the cN1A filament and identified important residues mediating polymerization. Expressing cN1A harboring mutations at these residues abolished assembly in cells, establishing that the cN1A filaments formed with purified protein *in vitro* are the basis for filaments we detected in cells and are likely to underlie cN1A accumulations in muscle tissue.

## Results

### cN1A localizes to the periphery of skeletal muscle fibers

cN1A is highly expressed in skeletal and cardiac muscle tissue,^13^ but its subcellular distribution in muscle fibers is not well understood. Tissue sections of skeletal muscle biopsies from healthy individuals (n=6) and IBM patients (n=6) were analyzed by immunohistochemistry (IHC) and immunofluorescence (IF). Using acidic ATPase staining as a reference for type 1 fibers,^27^ IHC showed that cN1A expression was distributed in a fiber type-specific checkerboard pattern in all biopsies analyzed, with the highest concentrations in type 2 fibers (Fig. 1A). Relatively strong cN1A staining was observed in all type 2 myofibers (healthy: 249/249, 100%; IBM: 246/247, 100%), but in only a minority of type 1 fibers (healthy: 8/513, 2%; IBM: 25/496, 5%). We confirmed the highest expression of cN1A was in type 2 fibers by IF (Fig. 1B). Staining of longitudinal muscle fiber sections, using antibodies to *α*-actinin as a marker for Z-disks, showed that cN1A localizes to disks flanking the Z-disks on both sides (Fig. 1C). Additionally, cN1A staining was most intense near the periphery of all muscle fibers analyzed by IF (Fig. 1D), where it appeared to assemble into a higher-order structure.

**Figure 1.**
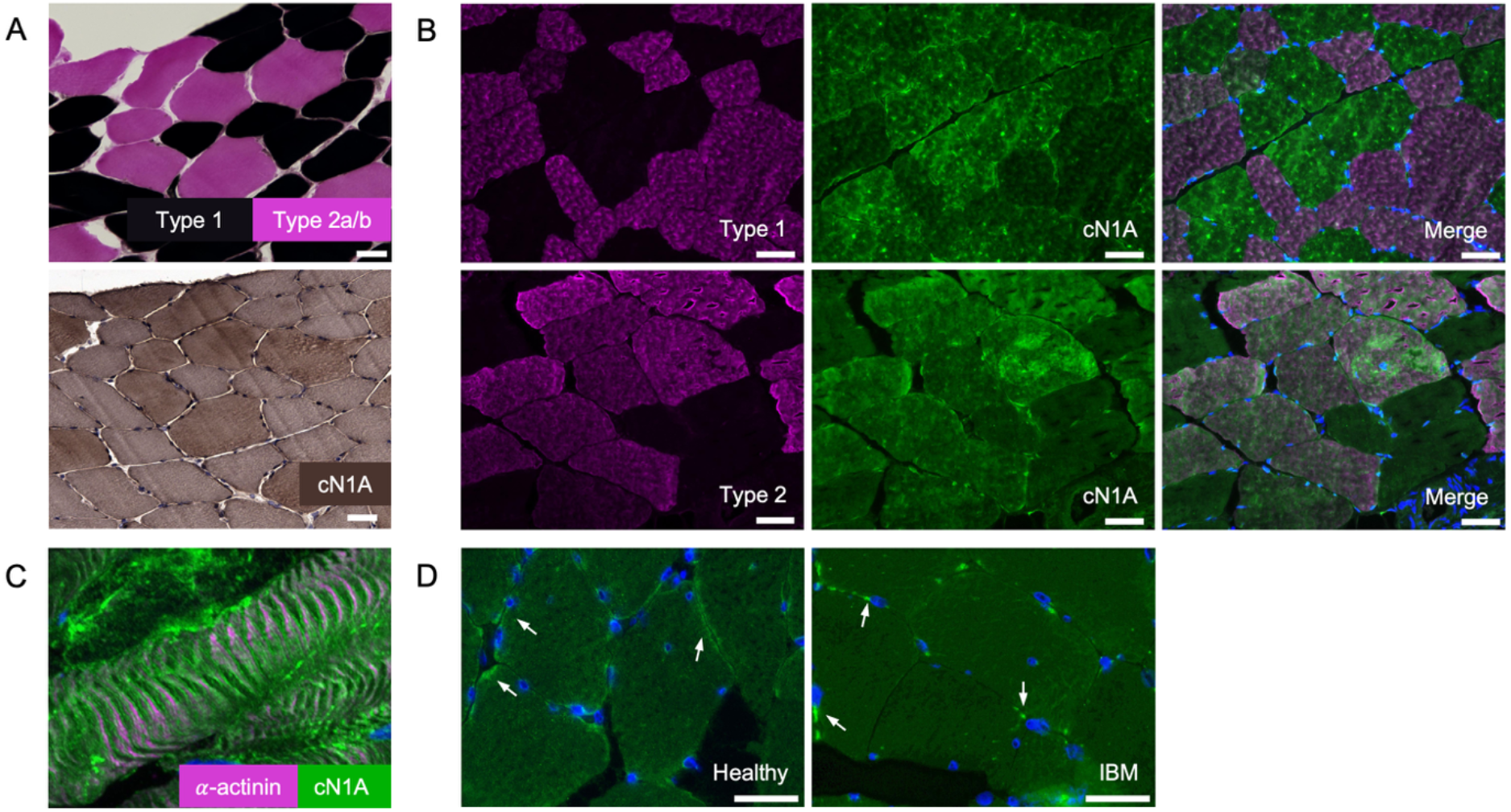
Expression and localization of cN1A in human skeletal muscle tissue. **A:** Serial healthy human skeletal muscle cross-sections were stained for ATPase activity at pH 4.2 (top) and with anti-cN1A antibody (bottom). **B:** Double immunofluorescence staining of healthy skeletal muscle cross-sections with a type 1 or type 2 fiber-specific antibody (magenta) and with anti-cN1A antibody (green). **C:** A longitudinal section of healthy skeletal muscle stained with antibodies to cN1A (green) and *α*-actinin (magenta) to visualize Z-disks. **D:** Skeletal muscle cross sections of healthy individuals and IBM patients were stained with anti-cN1A (green). Arrows point towards high-intensity cN1A staining near the periphery of the muscle fibers. Nuclei in panels B-D were stained with DAPI (blue). Scale bars: 50 µm (A-B) and 25 µm (D).

### cN1A assembles into micron-scale filaments in cultured cells

To determine if the cN1A filament assembly observed in HEp-2 cells in an earlier study^14^ was independent of cell type, we transfected HeLa and Hep-G2 cell lines under the same conditions. Using a CMV promoter, which drives heterogeneous, high protein expression (HE), both untagged cN1A (HE-cN1A) and cN1A fused to an N-terminal dimerization-deficient GFP^A206K^ variant^28^ (HE-GFP-cN1A) assembled into filamentous structures in all three cell lines, and remained similar in appearance at 24 or 48 h after transfection (Fig. 2A), indicating that cN1A filament formation is not specific to HEp-2 cells. cN1A filaments appeared mostly rod-like, but ring-shaped structures resembling IMPDH and CTPS filaments were also detected.^24,29^ To confirm that the GFP signal in the HE-GFP-cN1A-overexpressing cells was in fact labeling cN1A structures, HeLa cells transfected with HE-GFP-cN1A were stained with a cN1A-specific antibody, showing that GFP and cN1A signals indeed co-localize in the same structures (Fig. 2B).

**Figure 2.**
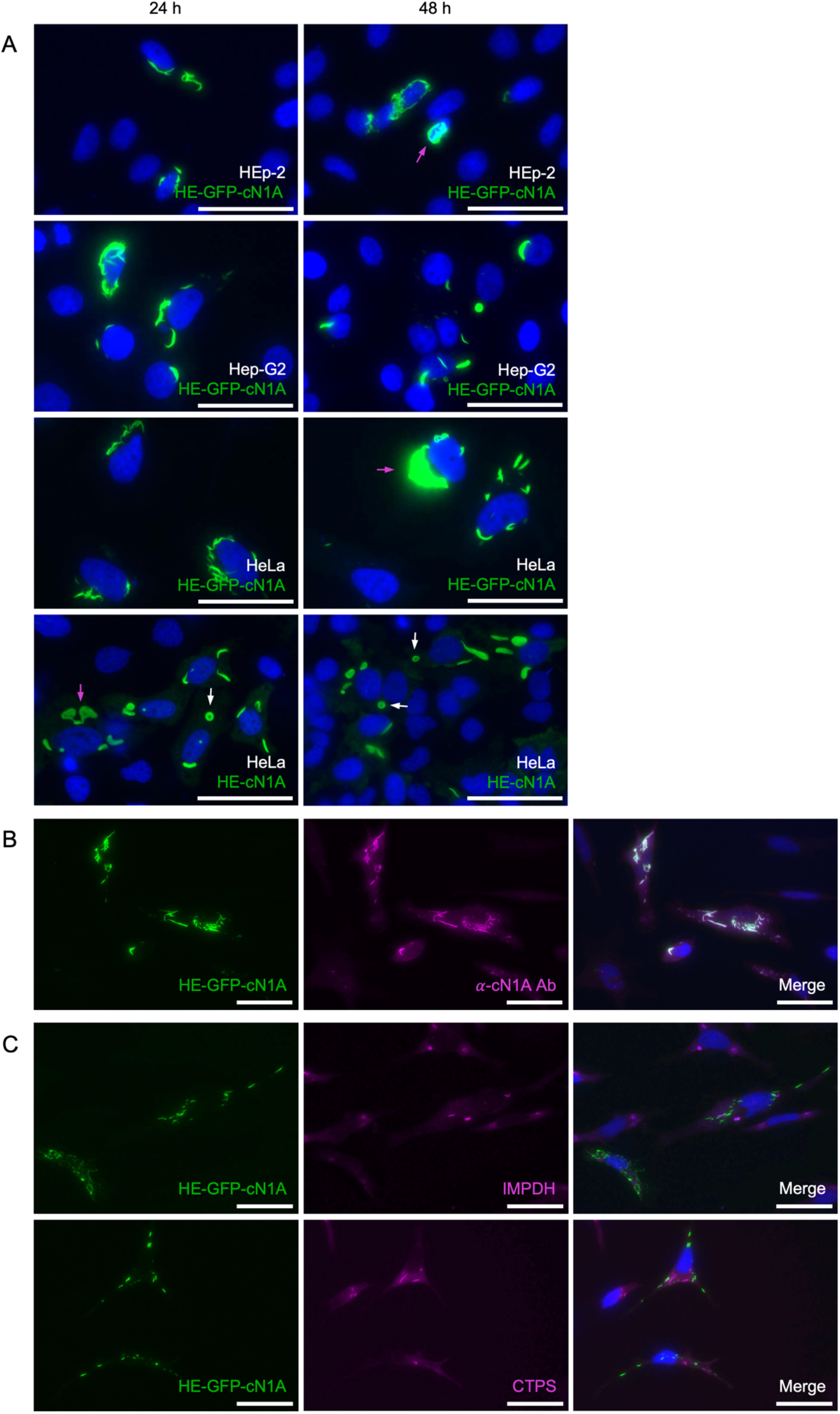
cN1A assembles into micron-scale filaments in cultured cells. **A:** Untagged or GFP-tagged cN1A (green) was expressed in HEp-2, Hep-G2, or HeLa cells using constructs resulting in high expression (HE) levels and imaged 24 or 48 h after transfection. Untagged cN1A (HE-cN1A) was visualized by immunofluorescence with anti-cN1A antibody and appeared similar in all three cell lines (HEp-2 and Hep-G2 cells not shown). White arrows point towards ring shapes. Magenta arrows point towards overexpression artifacts. **B:** HeLa cells transfected with a construct encoding GFP-cN1A (green) co-stained with polyclonal anti-cN1A antibody (magenta) 48 h after transfection. **C:** HeLa cells transfected with HE-GFP-cN1A (green) were treated with ribavirin (top) or DON (bottom) to induce formation of IMPDH or CTPS filaments (magenta), visualized by specific antibodies to these proteins. Nuclei in all panels were stained with DAPI (blue). Scale bars: 50 µm.

Since cN1A filaments resembled those of IMPDH and CTPS, human serum It2006 (anti-IMPDH)^24,26^ or an anti-CTPS1 antibody were used to assess co-localization with these enzymes. In HeLa cells transfected with the HE-GFP-cN1A construct and treated with ribavirin or DON to induce IMPDH or CTPS filament formation, respectively,^24^ cN1A filaments did not co-localize with either enzyme but assembled into distinct structures with similar morphology (Fig. 2C). Additionally, ribavirin or DON treatment did not seem to affect cN1A filament assembly; their morphology was comparable to filaments in untreated cells.

### cN1A forms filaments in cells even at low expression levels

Although cN1A filaments were distinct from IMPDH or CTPS filaments, we wanted to investigate if they were responsive to some yet unknown metabolic conditions in a similar manner. To do this, we wanted to establish cell culture conditions to study endogenous cN1A. Although cN1A mRNA has been detected in several human tissues,^13^ cN1A protein is detectable only in heart and skeletal muscle by western blotting.^30^ We readily detected cN1A protein in a western blot of human skeletal muscle lysate, but, as expected, it was barely detectable or undetectable in HeLa, HEp-2, or HEK293T lysate (Fig. 3A). cN1A was also undetectable by immunofluorescence in any of these cell lines (data not shown). Accordingly, we determined that transfection was necessary to continue investigating cN1A in cultured cells.

**Figure 3.**
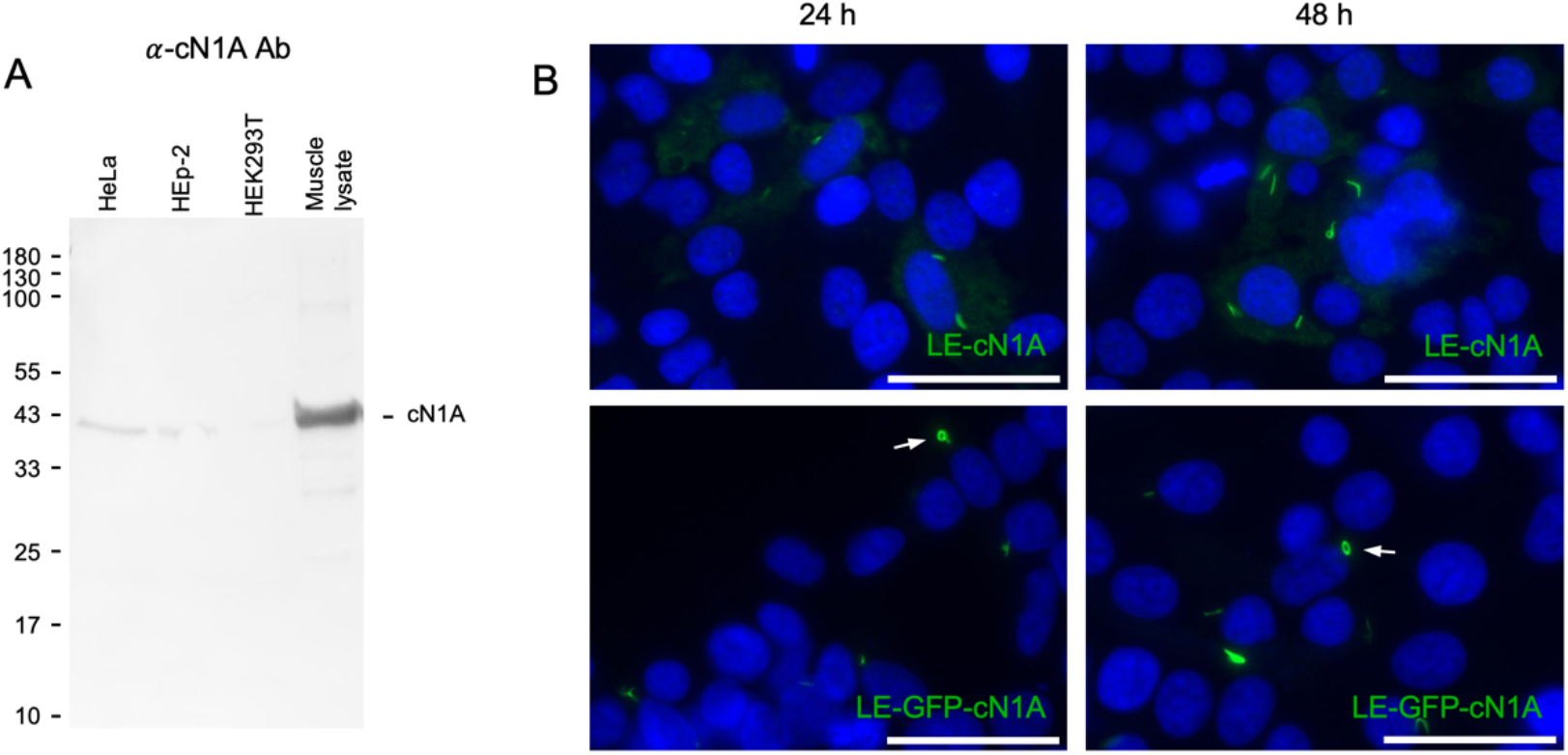
cN1A forms filaments in cells even at low expression levels. **A:** HeLa, HEp-2, and HEK293T cell lysates (7*10^5^ cells) and human skeletal muscle lysate (25 µg) were analyzed by western blotting using a polyclonal anti-cN1A antibody. The position of molecular weight markers is indicated on the left (in kDa). **B:** HeLa cells transfected with low expression LE-cN1A or LE-GFP-cN1A constructs visualized by anti-cN1A antibody or GFP signal 24 and 48 h after induction. Nuclei were stained with DAPI (blue). White arrows point towards ring shapes. Scale bars: 50 µm.

As expected, in our previous transfections using a CMV promoter, cN1A expression varied from cell to cell and was difficult to control by titrating plasmid concentration alone. Thus, we generated additional expression constructs using the Equalizer-L vector^31^ for low expression (LE) of untagged cN1A (LE-cN1A) and N-terminally GFP^A206K^-tagged cN1A (LE-GFP-cN1A). The Equalizer-L system uses both transcriptional negative feedback and posttranscriptional incoherent feedforward to produce constant gene expression and low cell-to-cell variability.^31^ Even at lowered expression levels, cN1A robustly assembled into filaments that were morphologically identical to the high expression condition (Fig. 3B), without the additional presence of overexpression aggregates we sometimes detected with expression driven by the CMV promoter (Fig. 2A, magenta arrows). With the LE constructs, cN1A expression levels were indeed reduced (lower signal in IF and undetectable by western blotting; results not shown) and less heterogeneous among individual cells. While this established that cN1A filament assembly is not an artifact of high overexpression, it did not provide us with overexpression conditions where we could conduct cell-based assays to investigate potential metabolic triggers for cN1A filament formation; its assembly is too stable.

### Purified cN1A spontaneously polymerizes *in vitro*

Since cN1A formed filaments regardless of expression level and without any apparent metabolic trigger, we wanted to determine if filament assembly is inherent to cN1A. We expressed and purified His_6_-tagged recombinant human cN1A from HEK293-6E cells (Supp. Fig. 1A) and imaged it by negative stain EM. Indeed, purified cN1A alone readily assembled into long filaments that were often hundreds of nanometers in length (Fig. 4A). We estimated the filaments to be just one monomer wide based on their diameter, suggesting unbounded, end-on-end stacking of cN1A monomers into filaments. Addition of substrates AMP or IMP,^32,33^ activators ADP or GTP,^13^ or the product adenosine did not affect filament formation (Supp. Fig. 1B).

**Figure 4.**
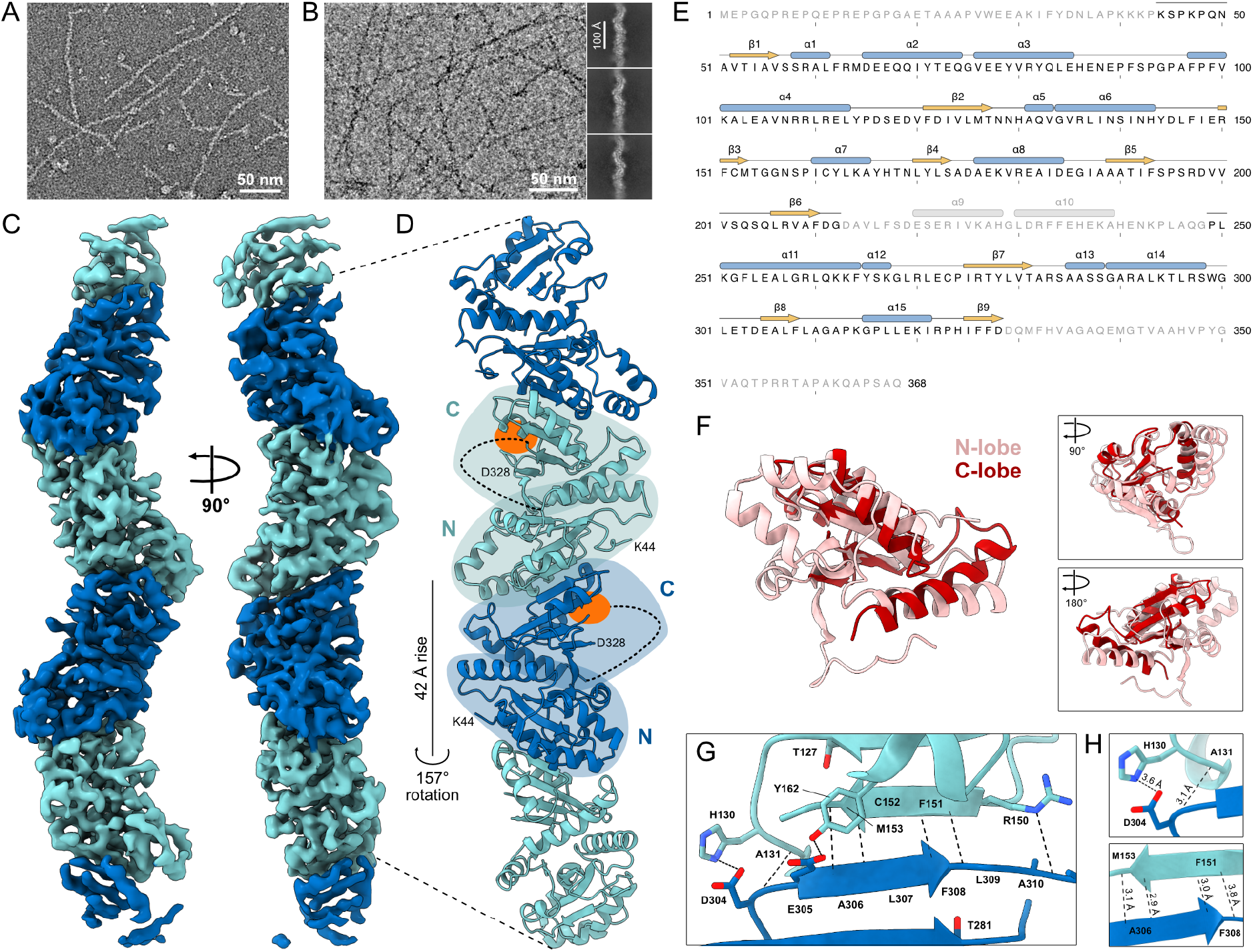
Cryo-EM structure of the cN1A filament. **A:** Negative stain EM of 1 µM cN1A in the absence of ligands (apo). **B:** Example cryo-EM micrograph of 30 µM apo cN1A low-pass filtered to 8 Å resolution (left) and an example subset of 2D classes of cN1A filaments (right). **C:** Cryo-EM volume containing four subunits (alternating teal and blue) of the cN1A filament at 4.1 Å resolution. **D:** Atomic model of four subunits of the cN1A filament. In the two central subunits, locations of the active sites (orange), most N-terminal (K44) and most C-terminal (D328) resolved residues, and missing residues 213-248 (dashed lines) are indicated. C: C-terminal lobe, N: N-terminal lobe. **E:** cN1A sequence with secondary structure shown (gold: β-strands, blue: helices) as annotated in our atomic model. Grey indicates unmodeled residues and predicted secondary structure based on pseudosymmetry and AlphaFold. The first helix in the AlphaFold model (residues 26-37) is omitted due to lower confidence in the prediction and lack of experimental support. **F:** Superposition of the N-lobe (pink) and C-lobe (red) from our model, calculated with TM-align. **G:** Magnified view of the filament assembly interface. Dashed lines indicate bonds calculated by PISA. **H:** Filament interface contacts selected for rational mutation design.

### Cryo-EM structure determination of the cN1A filament

To resolve the architecture of the cN1A filament, we collected a cryo-EM dataset in the absence of ligands. At 30 µM protein concentration in ice, filaments were one subunit wide and extended for hundreds of nanometers, similar to their appearance in negative stain at 30-fold lower protein concentration. They were not strictly linear, often arcing across micrographs, and 2D class averages showed a snaking filament that becomes blurred at the edges of the box, suggesting flexibility (Fig. 4B). We determined the filament structure at 4.1 Å resolution (Fig. 4C), sufficient to model backbone positions, revealing a polar filament built from end-to-end polymerization of cN1A monomers (Fig. 4D).

### Structural analysis of the cN1A monomer

No experimental structure has been reported for cN1A from any organism, although there are existing structures of related human nucleotidases. Our map resolves residues 44-212 and 249-328 of the 368-residue protein (Fig. 4E). The N-terminal 43 residues (1-43), the C-terminal 40 residues (329-368), and an internal 36-residue region (213-248) adjacent to the active site are not visible. The cN1A monomer comprises two structurally similar halves (“lobes”) with distant homology to the haloacid dehalogenase superfamily fold.^30^ Based on our structure, we used residues 44-204 as the N-terminal lobe (N-lobe) and 205-328 as the C-terminal lobe (C-lobe). The two lobes are pseudosymmetric, which is apparent when the two lobes are aligned based on structure with TM-align^34^ (TM-score: 0.68, RMSD: 2.9 Å), despite the unmodeled residues 213-248 (Fig. 4F). The lobes are related by a rotation of 178.8° and translation of 1.8 Å but share only 17% sequence identity (14 identical residues in 84 aligned positions), indicating an internal duplication in which the fold has been retained while the sequence relationship has been erased. Aligning the two lobes from the AlphaFold predicted structure, with residues 213-248 intact but residues 1-43 and 329-368 trimmed to match our model, supports that the two lobes are in essentially identical folds (Supp. Fig. 3A; TM-score: 0.85, RMSD: 2.1 Å), with the inclusion of residues 213-248 only raising the sequence identity to 21% over 122 aligned positions. AlphaFold predicts that both N-and C-termini are mostly unstructured and flexible, while residues 213-248 are folded, suggesting this domain is moving enough that we could not resolve it by cryo-EM.

Structure-based sequence alignments of the N- and C-lobes generated by TM-align for our experimentally determined structure (Supp. Fig. 3B) and the AlphaFold predicted structures (Supp. Fig. 3C) agree at 77 out of the 84 aligned positions, with remaining positions differing by single-residue shifts adjacent to gaps.

While their folds remained the same, the functions of the lobes have diverged. Three catalytic motifs have been identified in the intracellular nucleotidases.^30,35^ For cN1A, all three are located in the C-lobe: motif I (211-216), motif S (244-254), and motif III (320-329). The catalytic nucleophile D211 sits at the first position of the DXDX[T/V][L/V/I] catalytic motif I within the sequence DGDAVL, while the N-lobe carries the sequence SSR-AL in the equivalent position, maintaining only the end leucine of the motif (Supp. Fig. 3C). Motif S, P(X)_7-8_[R/K]XGF[W/L], which forms hydrogen bonds with the substrate base, is PLAQGPLKGFL in the C-lobe and PFSPGPAFPFV in the N-lobe, again losing critical motif residues. Motif III, K(X)_5_D(X)_3_DD, required to form hydrogen bonds with AMP and Mg^2+^, is KIRPHIFFDD in the C-lobe and AYHTNLYLSA in the N-lobe, losing the important lysine and aspartate residues. The N-lobe is therefore a degenerate pseudo-active site that preserves the catalytic fold but not the chemistry.

### Architecture of the cN1A filament assembly interface

Monomers stack end-to-end into a single-stranded filament with a rise of 42.4 Å and a twist of 157.3° per subunit (calculated at the modeled interface), corresponding to 2.3 subunits per turn. The filament is one subunit in cross-section and is polar: each interface buries the N-lobe face of one subunit against the C-lobe face of the next (Fig. 4D). Analysis of the interface with PISA^36^ reveals the interface buries 1160 Å^2^ per subunit, 8.8% of the subunit surface, and is strictly segregated by lobe: all 33 contributing residues of one subunit lie in the N-lobe (45-167) and all 33 of its partner subunit in the C-lobe (207-321).

The largest contribution to the interface comes from the creation of a β-sheet: a β-strand on the edge of the N-lobe (150-153) pairs antiparallel with a strand on the edge of the C-lobe of the adjacent subunit (305-308) (Fig. 4G). Each β-strand has a single intra-chain neighboring threonine, T127 and T281, respectively, themselves a pseudosymmetric equivalent pair (Supp. Fig. 3B). In our model, three inter-subunit backbone hydrogen bonds define the interaction: F151 O to F308 N (3.0 Å), M153 N to A306 O (2.9 Å), and M153 O to A306 N (3.1 Å). The inter-subunit β-sheet is continuous through the filament core: a β-sheet extends through the N- and C-lobes of each cN1A monomer itself, and is then further extended to the next subunit during polymerization, similar to β-strand addition described for other protein complexes.^37^ Peripheral polar contacts flank this β-sheet, including Y162 OH to E305 OE2 (2.9 Å), R150 NE to A310 O (2.9 Å), H130 ND1 to D304 OD1 (3.6 Å), and E77 OE1 to R270 NH2 (3.9 Å), although side chain placement was not always obvious at 4.1 Å resolution. F151 and F308 side chains may also pack against one another across the interface (3.8 Å between side chain heavy atoms), but F151 could not be placed with sufficient confidence to define this contact.

The active site is not occluded by filament formation. The nucleophile D211 is 12.8 Å from the nearest atom of the adjacent subunit (D211 OD1 to F151 O), and the attachment points of residues 213-248 (not visible) project away from the interface. Of the residues implicated in AMP binding, only D211 is resolved; D213 and V215 fall within the unmodeled region.

### Structure-guided mutations abolish cN1A filament assembly in cells

Next, we wanted to test if the specific contacts we identified at the filament assembly interface *in vitro* (Fig. 4H) are required for filament assembly in cells. We transfected HeLa cells with LE constructs encoding GFP-tagged WT, A131E, H130E-A131E, and F151K-F308K cN1A variants (Fig. 5A). The single substitution A131E, intended to introduce electrostatic repulsion against D304 and E305 of the adjacent subunit, did not detectably affect filament formation compared to WT.

**Figure 5.**
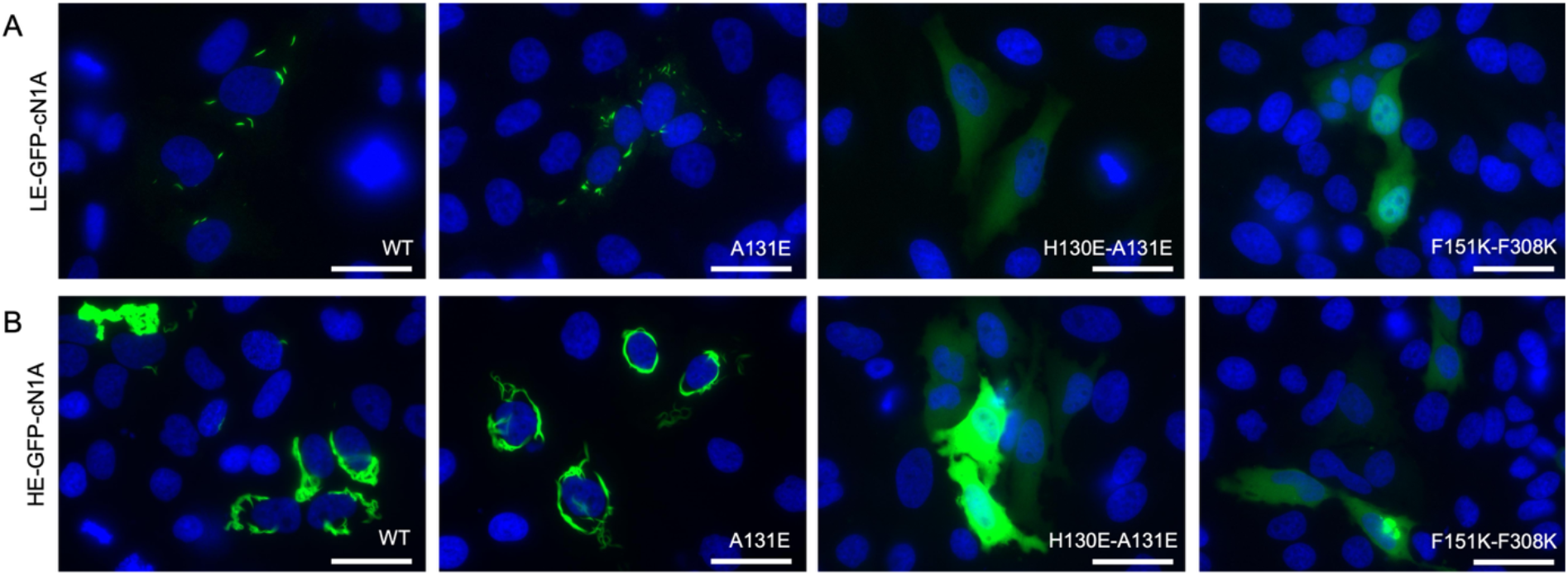
Structure-guided mutations disrupt assembly of cN1A filaments in cells. **A:** HeLa cells transfected with LE constructs expressing GFP-tagged wild type (WT) or A131E, H130E-A131E, and F151K-F308K cN1A variants imaged 24 h post-induction. **B**: HeLa cells transfected with HE constructs expressing GFP-tagged WT or the same variants imaged 24 h post-transfection. Nuclei were stained with DAPI (blue). Scale bars: 50 µm.

However, two double mutations abolished filament assembly outright, producing only diffuse cytosolic staining: H130E-A131E, adding additional charge next to A131 in place of H130, whose side chain directly contacts D304, and F151K-F308K, putting two charged residues across from each other in positions typically forming a backbone hydrogen bond (Fig. 4H). Both mutants target residues whose side chains are >90% buried on assembly. We then transfected cells with HE constructs to determine if the weakened assembly interface interactions could be overcome by protein concentration. Even at high overexpression levels, H130E-A131E and F151K-F308K mutants could not form filaments (Fig. 5B). Disrupting either the H130-D304 contact or the F151-F308 backbone pairing is sufficient to abolish filament formation in cells, confirming that the interface we identified is the primary driver of cN1A assembly.

## Discussion

Since the identification of anti-cN1A autoantibodies as a biomarker for IBM just over a decade ago,^3,4^ mounting evidence suggests that cN1A protein accumulates into higher-order structures in (patient) muscle tissue, especially near the nucleus,^3,7,11,12^ rimmed vacuoles,^3,11,12^ sarcoplasmic membranes, and in the sarcoplasm.^7^ Previous research has also associated cN1A with contractile elements in cardiac muscle, including a striated cN1A staining pattern in pigeon heart tissue^38^ and association with myofilaments in dog heart.^39^ In this study, we found increased cN1A expression in type 2 compared to type 1 myofibers and cN1A staining flanking the Z-disks of sarcomeres in healthy human skeletal muscle tissue. During muscle contraction, the myosin heads hydrolyze ATP while moving towards the Z-disks, meaning cN1A appears to be located at regions where relatively high concentrations of ADP are produced. ADP is subsequently reconverted to ATP in the mitochondria by the action of myokinase, which converts 2 ADP molecules into ATP and AMP, the latter of which may be converted by AMP deaminase into IMP or by cN1A into adenosine in the complicated network of salvage pathways that maintain balance of nucleotide pools.^40^ Type 2 fibers mediate short, intensive contractions, associated with massive ATP hydrolysis, leading to an increase in ADP and AMP levels.^41^ Since ADP strongly activates cN1A *in vitro*, cN1A is likely activated under these circumstances.^32^ When these conditions are prolonged, the pH drops, thereby inhibiting cN1A.^42,43^

In addition to its localization in sarcomeres, we found cN1A accumulations close to nuclei and in the periphery of both healthy and IBM myofibers, similar to previous reports.^3,7,11,12^ This assembly of endogenous cN1A in tissues was paralleled by robust filament assembly of ectopically expressed cN1A in cultured human cell lines. The mechanism of filament assembly was explained by resolving its structure: cN1A polymerizes head-to-tail through binding between cN1A monomers and requires no ligand. An identical binding site becomes available with every subunit added, so polymerization is limited only by the concentration of free monomers.

Each cN1A lobe shows high structural similarity to other human nucleotidases, especially the cytosolic and mitochondrial 5’(3’)-deoxyribonucleotidases cdN (gene: NT5C) and mdN (gene: NT5M), which share an identical fold and also contain catalytic motifs I, S, and III.^30,35,44^ The core fold of cN1A closely follows that of cdN (PDB: 4L57)^44^ and mdN (PDB: 1MH9)^35^ with a few minor differences (Supp. Fig. 4). The “small domain” of cdN/mdN forms a truncated four-helix bundle^35^ that is shortened to a helix-turn-helix in cN1A of approximately half the total residues (21 vs. 47 residues). This corresponds to residues 65-85 in the N-lobe and residues 219-239 of the C-lobe, part of the unresolved region in our cryo-EM map, indicating freedom of movement of this folded small domain in the C-lobe. cN1A retains one helix (residues 176-185) of the canonical Rossmann fold that is lost in cdN/mdN, which have a loop in that position instead (cdN: 145-158; mdN: 176-189);^35^ the equivalent helix in the C-lobe (residues 331-339 in the AlphaFold prediction) is not visible in our map. In place of a short helix seen in cdN/mdN, the cN1A fold also incorporates an unstructured loop positioned just before the β-strands at residues 150-153 and 305-308, potentially facilitating formation of the filament assembly interface. Helices positioned here would probably prevent a hydrogen bond between S284 OG and E149 OE2 (3.0 Å), and might alter the positioning of D304, but it is unclear if other filament assembly contacts would be disrupted or if different stabilizing bonds from the helices would equally facilitate filament formation.

While our cryo-EM structure describes a single-stranded filament ∼6 nm in diameter, subcellular cN1A filaments visible by fluorescence microscopy are much thicker, implying the subcellular structures are actually bundles of individual filaments, as is the case with IMPDH filaments.^45–47^ Because we did not observe bundles *in vitro*, any lateral contacts needed for their assembly are either impeded by the C-terminal His_6_ tag or require an unknown cellular factor or post-translational modification that was not preserved in our purified protein. Nevertheless, it is intriguing to speculate that the unmodeled N- and C-termini (43 and 40 residues long), predicted to be relatively unstructured and flexible, could be involved. Both residues 44 and 328 point out from the surface of the filament, suggesting the termini could extend well beyond the 3 nm filament radius, even at modest extension, and could therefore reach an adjacent filament or bind an interactor that tethers two filaments. We know for certain that fusion of GFP to the N-terminus of cN1A does not interfere with filament formation in cells, but future experiments could determine if measurable differences in subcellular filament width arise from altering each terminus separately. It is also possible that, rather than interactions of the termini, more rigid interactions between individual cN1A filaments drive formation of higher-order bundles, which may curve into bent rod-like shapes, reminiscent of structures formed by budding yeast CTPS filaments.^48^ The unresolved C-lobe small domain (residues 213-248), which appears to move freely even when the filament is assembled, might facilitate such an interaction. However, that our interface mutations abolish filament assembly entirely indicates that the β-sheet contact, not lateral association, is the primary driver of filament formation, and bundling might be a secondary organization of pre-formed filaments. Imaging of cells containing cN1A filaments by TEM could resolve this directly and may also provide additional insight into what extent cN1A filament assembly contributes to cN1A accumulation in skeletal muscle, which may be larger extensions of subcellular bundles.

Subcellular cN1A filaments resemble those formed by other nucleotide metabolic enzymes, but the resemblance is morphological rather than mechanistic. Inhibition of *de novo* nucleotide synthesis in cultured cells induces rapid, reversible assembly of IMPDH and CTPS filaments, which disassemble on return to homeostasis,^19,24,49–52^ and both are dynamically regulated *in vivo* during lymphocyte proliferation.^17,18,20^ In both cases, assembly is controlled by allosteric ligand binding and constitutes an additional layer of regulation on enzymes whose activity must be tightly controlled. Assembly of cN1A appears to be constitutive rather than regulated, with the filament as the default oligomeric state. Filaments formed in every cell line we tested, regardless of expression level or fusion to GFP, and neither substrates (AMP, IMP), activators (ADP, GTP), nor the product adenosine affected polymerization of the recombinant protein. Our work suggests that ligand binding does not control filament assembly in the way it does for IMPDH and CTPS filaments, but does not rule out minor rearrangements of the assembly interface or conformational changes to accommodate ligand binding that we cannot detect at the resolution of negative stain EM. The robustness of assembly makes it difficult to determine subcellular conditions that regulate cN1A filaments and may require a muscle tissue culture model to identify tissue-specific regulatory factors.

While it remains unclear if filament assembly is regulated, our structure suggests that cN1A is catalytically active as a filament, though direct measurement of the activity of assembled versus monomeric cN1A would be required to confirm this. The active site faces away from the assembly interface and is neither buried nor occluded in the filament. Although we were unable to model D213 and V215 of catalytic motif I^30,35^ that lie within the unresolved 213-248 segment, we hypothesize that substrate binding positions this region adjacent to the active site, which predicts that a structure determined with AMP would resolve it. AlphaFold predicts the segment to adopt a fold containing a helix-turn-helix that closely resembles residues 59-94, the pseudosymmetric equivalent region of the N-lobe. The two lobes of cN1A are related by a near-exact internal two-fold symmetry, yet only the C-lobe retains the catalytic residues, making the N-lobe a degenerate pseudo-active site. While it lost its catalytic function, the N-lobe might have been repurposed as a polymerization surface. Although probably not a strong contributor to the filament assembly interface, at least in the apo-cN1A filament, our model indicates a potential interaction between E77 (in the N-lobe equivalent of the unresolved C-lobe segment) and R270.

Recently, we showed that anti-cN1A autoantibodies in the sera of IBM patients can bind cN1A filaments in cultured cells.^14^ This resulted in the development of a cell-based assay for the detection of anti-cN1A autoantibodies, which showed a higher sensitivity compared to existing ELISA assays, suggesting that cN1A filaments contain specific autoepitopes that are lacking in immobilized, purified cN1A peptides. However, our structure adds an unexpected observation. All three mapped autoantibody epitopes (residues 25-47, 221-243, and 345-367)^4^ lie in regions our cryo-EM map does not resolve and are unmodeled. In an ELISA with the full-length recombinant protein, these segments would remain disordered or flexible whether or not the protein is assembled into a filament, making it unlikely that increased accessibility of one of these epitopes explains the difference in assay sensitivity. However, the formation of more stable conformations by these epitopes inside cells, either by lateral filament contacts or by substrate binding, may lead to more efficient recognition of these epitopes in our cell-based anti-cN1A assay. Alternatively, such interactions may also lead to the formation of neo-epitopes that are recognized by a separate class of anti-cN1A autoantibodies, likewise increasing sensitivity.

In hepatitis C patients, treatment with interferon-α and ribavirin correlates with an anti-IMPDH (anti-rods/rings) autoantibody response.^25,53^ One hypothesis is that ribavirin-induced IMPDH filament formation, combined with proinflammatory interferon-α and the immunostimulatory environment of hepatitis C viral infection, drives loss of tolerance.^25^ Our observations that cN1A spontaneously and constitutively forms filaments suggest that loss of immune tolerance is not related to its oligomerization. Unlike IMPDH, there is no trigger for filament assembly, like ribavirin treatment, that coincides with an inflammatory trigger. It is also important to note that cN1A accumulations are observed in myofibers from both IBM patients and healthy individuals. Nevertheless, our data do not exclude the possibility that alterations in localization or excessive accumulation of cN1A in myofibers contribute to the initial immune response in IBM. Additionally, aberrant release of cN1A filaments from myofibers could contribute to loss of immunological tolerance, prior to onset or during early development of IBM.

Recently, we identified three monomethylation sites in cN1A isolated from human skeletal muscle tissue.^54^ Mutation of these residues did not appear to affect cellular filament assembly. Interestingly, two of these methylation sites reside in the previously mapped epitope (221-243) within the unresolved C-lobe small domain. An ELISA analysis with a synthetic methylated cN1A peptide demonstrated increased anti-cN1A autoantibody reactivity, consistent with the possibility that the methylations structurally stabilize the epitope in a manner that supports its recognition by the paratope. Additional studies are required to assess if and how these posttranslational modifications are involved in activation of the immune system in IBM patients.

Here, we report the inherent propensity of human cN1A to polymerize into filaments. Our cryo-EM structure of the filament provides a molecular basis for further investigation into the physiological role of cN1A in nucleotide metabolism, its localization and regulation in myofibers, and its potential contribution to the development or progression of IBM as an autoantigen.

## Materials and Methods

### Patients and muscle samples

A cross-sectional study was conducted using muscle samples from IBM patients and healthy controls. Patients with IBM fulfilling the 2010 modified Griggs criteria^55^ for definite or possible IBM were recruited from the Radboud University Medical Center (Nijmegen, the Netherlands). Healthy individuals without a history of neuromuscular disease were recruited as controls. Exclusion criteria for all participants were as follows: age <18 or ≥65 years, diabetes mellitus, chronic obstructive pulmonary disease, chronic heart failure, current malignancy, previous treatment with chemotherapy and/or radiation therapy, use of corticosteroids for more than two weeks in the past 5 years, current use of statins, or being wheelchair bound. The Medical Ethics Review Committee region Arnhem-Nijmegen approved the use of muscle biopsies for this study (no. 2011/181).

### Immunohistochemical (IHC) and immunofluorescent (IF) staining of muscle tissue

Needle muscle biopsies were taken from the musculus vastus lateralis or the tibialis anterior, frozen in isopentane, and stored at −80 °C. Serial cryostat sections (8-10 µm) were taken from the muscle tissue and stored at −20 °C. ATPase (pH 4.2) analysis was carried out in accordance with protocols described by Dubowitz and coworkers.^56^ Sections were thawed and air-dried for 30 min, fixed in cold acetone (4 °C) for 12 min, and then pre-incubated with Tris-buffered saline (TBS) containing 0.1% Tween-20, 1% BSA, and 20% goat serum for 30 min at room temperature (RT) or 1% BSA/PBS supplemented with 10% normal sheep serum (NSS) for 1 h. After blocking, excess fluid was removed, and primary antibody was added to tissue sections and incubated overnight at 4 °C (IHC: rabbit anti-cN1A, 1:400 in 1% BSA/TBS, Ab75101, Abcam; IF: rabbit anti-cN1A, 1:50 in 1% BSA/TBS, HPA050283, Atlas Antibodies). Incubations with other primary antibodies were performed in 0.1% BSA/TBS for 1 h at RT: mouse anti-myosin heavy chain type 1 (BA-D5, DSHB), mouse anti-myosin heavy chain type 2 (A4-74, DSHB), or mouse anti-*α*-actinin (A7811, Sigma-Aldrich). Subsequently, sections were washed with TBS (3 x 5 min) and incubated with secondary antibody (IHC: biotinylated donkey anti-rabbit IgG, 711-065-152, 1:200 in 1% BSA/TBS, Jackson ImmunoResearch; IF: goat anti-rabbit IgG Alexa Fluor 555, 1:500 in 1% BSA/TBS, A11036, Invitrogen, or goat anti-mouse IgG Alexa Fluor 568, 1:500 in 1% BSA/TBS, A11004, Life Technologies) for 1 h at RT. After washing with TBS (3 x 5 min), nuclei were stained for 1 min with 1 µg/mL DAPI in TBS, followed by washing with TBS. Stained tissue sections were embedded in Mowiol mounting medium and stored overnight at 4 °C. For IHC, an enhancement step was performed using the Vectastain ABC kit (PK-4000, Vector Laboratories) as described by the manufacturer. Following a final TBS wash, antibody binding was visualized by 3,3’-diaminobenzidine-H_2_O_2_ chromogen (DAB) precipitation for 10 min in the dark, after which the staining reaction was stopped with running tap water (10 min). DAB staining was intensified with 1% copper sulfate in TBS for 5 min and again stopped with running tap water for 10 min before staining nuclei with hematoxylin for 20 min. After a final wash with water for 15 min to stop hematoxylin staining, sections were dehydrated (three consecutive 5 min washes with 50, 70, 96, and 100% alcohol and three 5 min washes with xylene) and embedded in Quick-D mounting medium. All incubation steps were performed in a humid chamber in the dark. Negative control sections underwent the same staining protocol, but incubation with primary antibody (anti-cN1A) was omitted.

### Histological assessment

Semi-quantitative analysis was performed by selecting an area of interest of 100 to 150 good-quality myofibers in sections stained with Hematoxylin-phloxine (H-phlox) and analyzing the corresponding region in images of other staining. Light micrographs were taken using VisionTek (99990060-0512, SakuraTM) and Phillips Intellisite Pathology Solution scanners (452220700811, DPS). Quality requirements for the inclusion of muscle sections were the presence of an intact area of muscle in which the myofibers were closely knit in all stains and the possibility to compare them in serial sections. The number of myofibers was quantified for fiber types and the proportion of myofibers with high-intensity cN1A staining was assessed.

### DNA constructs

We generated the reference sequence of cN1A (NM_032526.1) by substituting a single nucleotide in an existing plasmid containing full-length cDNA for cN1A.^4^ We then subcloned the cN1A reference sequence into pcDNA5/FRT/TO and Equalizer-L vectors (Addgene plasmid #169732) for fluorescence imaging experiments.^31^ The latter plasmid allows low protein expression in an inducible system. In both plasmids, the dimerization-deficient GFP^A206K^ variant^28^ was inserted in frame at the N-terminus of the cN1A coding sequence, with a linker (YSDLEENLYFQGSFT). For purification of recombinant protein from HEK293-6E cells, the cN1A coding sequence was inserted into the pIRES-EGFP-puro vector (Addgene: #45567), yielding the pIRES-cN1A-His_6_-GFP plasmid, which allows the production of C-terminally His_6_-tagged cN1A. IRES-mediated GFP expression occurs in parallel to facilitate the differentiation between transfected and non-transfected cells. Mutations in the pcDNA5/FRT/TO-GFP^A206K^-cN1A and Equalizer-L-GFP^A206K^-cN1A constructs were introduced with a PCR-based site directed mutagenesis.

### Cell culture

Adherent HEp-2, Hep-G2, HeLa, and HEK293T cell lines were maintained between 40 and 80% confluency. Cells were cultured in DMEM high glucose with GlutaMAX and pyruvate (Gibco), 10% fetal bovine serum (FBS) (Merck), and 10 U/mL penicillin-streptomycin (Gibco). The suspension cell line HEK293-6E was passaged twice a week to maintain a concentration under 2 million cells per mL in Freestyle F17 medium supplemented with 10 U/mL penicillin-streptomycin (Gibco), 0.1% Kolliphor P188, and GlutaMAX. All cells were grown at 37 °C in a humidified incubator with 5% (v/v) CO_2_. Suspension cells were cultured shaking at 270 rpm.

### cN1A transfection and induction of IMPDH/CTPS filaments in cell culture

HEp-2, Hep-G2, and HeLa cells were seeded on coverslips coated with 0.1 mg/mL poly-L-lysine in a 24-well plate or in an 8-well µ-Slide with IbiTreat (80806, Ibidi), then transfected with pcDNA5/FRT/TO or Equalizer-L plasmids encoding cN1A (indicated in the text and figure legends) the following day using Lipofectamine 2000 (Invitrogen). The transfection mixture was removed and fresh medium was added to the wells 7 h after transfection. To induce protein expression in cells transfected with Equalizer-L plasmids, the fresh medium contained 1 µg/mL doxycycline. To induce the formation of IMPDH or CTPS filaments, the fresh medium also contained 1 µM ribavirin or 1 mM 6-diazo-5-oxo-L-norleucine (DON), respectively.

### Immunofluorescence of transfected, cultured cells

At 24 or 48 h, the cells were washed twice with PBS before fixation with 4% paraformaldehyde in PBS for 10-20 min at RT. The cells were washed twice with 5% BSA in PBS between every staining or permeabilization step. Cells were permeabilized with 0.1% Triton X-100 in PBS for 20 min and blocked with 5% BSA in PBS for 1 h at RT. All antibodies were diluted in 5% BSA in PBS and cells were incubated with antibodies for 1 h at RT. cN1A was stained with rabbit anti-cN1A (10 µg/mL, ab190214, Abcam) or rabbit anti-cN1A (2 µg/mL, HPA050283, Atlas) and goat anti-rabbit IgG Alexa Fluor 647 (4 µg/mL, A21244, Invitrogen) or goat anti-rabbit IgG Alexa Fluor 647 (2 µg/mL, A27040, Invitrogen). IMPDH was stained with human serum It2006 (1:500, kindly provided by Edward Chan)^26^ and anti-human-Ig Alexa Fluor 647 (1 µg/mL, 2010-31, Southern Biotech). CTPS was stained with goat anti-CTPS1 (0.4 µg/mL, sc-131474, Santa Cruz) and donkey anti-goat IgG Alexa Fluor 568 (2 µg/mL, A11057, Life Technologies). After the secondary antibody was added, all incubation steps were performed in the dark. To stain nuclei, cells were incubated with 1 µg/mL DAPI in PBS with 5% BSA and incubated for 10 min. Samples were washed twice with PBS before mounting the coverslip on a histology slide with mounting medium (Dako). Samples were kept in the dark at 4 °C until imaging.

### Western blotting

Extracts from non-transfected HeLa, HEp-2, or HEK293T cells (7*10^5^ cells) and human skeletal muscle lysate (25 µg) were subjected to 12% polyacrylamide gel electrophoresis. A pre-stained protein marker (26619, Thermo Fisher) was used for size determination. Separated proteins were transferred to a nitrocellulose membrane (Amersham Protran) by electroblotting. The blot was blocked with 5% non-fat dry milk in PBS containing 0.05% Tween-20 (PBST). cN1A was detected by subsequent incubations with rabbit anti-cN1A (0.5 µg/mL, A30631, Boster) and goat anti-rabbit IgG IRDye680 (0.2 µg/mL, 926-68071, LI-COR Biosciences). GFP was detected by mouse anti-GFP (0.2 µg/mL, F56-6A1.2.3, Invitrogen) and goat anti-mouse IgG IRDye800 (0.1 µg/mL, 926-32210, LI-COR Biosciences). Bound antibodies were visualized using the G:Box chemi XX6 (Genesys).

### Production of recombinant cN1A-His_6_

Recombinant C-terminally His_6_-tagged cN1A protein (cN1A-His_6_) was produced in HEK293-6E suspension cells. One million cells per mL were transfected with 0.5 µg/mL DNA and 0.75 µL/mL FectoPRO (Polyplus). Thirty minutes later, 0.5 µL/mL Booster was added. After 4 days, the cells were collected by centrifugation (10 min at 1,000xg) and washed once with PBS. Cells were lysed in lysis buffer (50 mM sodium phosphate, pH 8.0, 300 mM sodium chloride, 2% Tween-20, 50 mM imidazole, 10% glycerol, 1x Complete protease inhibitor (Roche)) and sonicated for 10 cycles, 15 sec on, 15 sec off. To remove insoluble material, the lysate was centrifuged for 10 min at 20,000xg at 4 °C. cN1A-His_6_ was purified from the lysate by nickel affinity chromatography. Ni-NTA agarose beads (Qiagen) were washed with column buffer (50 mM sodium phosphate, pH 8.0, 300 mM sodium chloride, 20 mM imidazole) and incubated with the cell lysate supernatant overnight at 4 °C while rotating head-over-head. The next day, the beads were allowed to form a column, which was washed three times with 3 column volumes of column buffer supplemented with 10% glycerol. Bound cN1A-His_6_ protein was eluted with elution buffer (50 mM sodium phosphate, pH 8.0, 300 mM sodium chloride, 500 mM imidazole, 10% glycerol). The buffer was exchanged using a 7K MWCO Zeba Spin Desalting Column and concentrated using a 3 kDa MWCO Millipore spin filter. The final storage buffer was 50 mM sodium phosphate, pH 8.0, 300 mM sodium chloride, 10% glycerol, and 1 mM DTT. The concentration of recombinant cN1A-His_6_ was determined with a BCA assay. The recombinant protein was aliquoted, flash-frozen, and stored at −80°C.

### Negative stain electron microscopy

Protein in storage buffer was thawed and diluted to 1 or 2 µM in buffer containing 50 mM sodium phosphate (pH 7.4), 150 mM sodium chloride, and 1 mM DTT (final imaging buffer: 50 mM sodium phosphate, 154 mM sodium chloride, 0.26% glycerol, 1 mM DTT). Where indicated, adenosine, AMP, ADP, IMP, or GTP were added to 10 mM final concentration in addition to 20 mM magnesium chloride. Samples were incubated for 15 min at RT before 3 µL of sample was applied to glow-discharged, carbon-coated grids and incubated for 1 min. Samples were blotted, and grids were washed in water prior to staining with 0.7% uranyl formate. Samples were imaged on an FEI Tecnai T12 TEM operating at 120 kV with a Gatan Ultrascan 4000 CCD camera controlled by DigitalMicrograph software. Micrographs were collected at a nominal 67,000X magnification with a 1.6 Å pixel size.

### Cryo-EM sample preparation and data collection

Protein in storage buffer was thawed and diluted to 30 µM in buffer containing 50 mM sodium phosphate (pH 7.4), 150 mM sodium chloride, 1 mM DTT, and fluorinated octyl maltoside (FOM; O310F, Anatrace) added to 0.05% concentration (final imaging buffer: 50 mM sodium phosphate, 209 mM sodium chloride, 3.9% glycerol, 1 mM DTT, 0.05% FOM). After incubation for 15 min at RT, 3 µL of sample was applied to a glow-discharged C-Flat 300 mesh 1.2/1.3 holey carbon grid (Electron Microscopy Sciences), blotted at 22 °C with 100% relative humidity, and plunge-frozen in liquid ethane with a Thermo Scientific Vitrobot Mark IV vitrification system. High-throughput data acquisition was performed with Leginon software^57^ controlling a Thermo Scientific Titan Krios G3 TEM at 300 kV equipped with a Gatan image filter and K3 Summit direct electron detector. A total of 3,056 movies were collected in counting mode with a pixel size of 0.835 Å and dose per frame of 0.49 e-/Å^2^. Movies were captured in 100 frames with a total exposure dose of 49 e-/Å^2^.

### Cryo-EM data processing

Our cryo-EM data processing workflow is summarized in Supp. Fig. 2. All data processing was performed in cryoSPARC v4.7.^58^ Movies were aligned, corrected for beam-induced motion, and dose-weighted with patch motion correction. Micrographs with CTF fit resolution above 6 Å from patch CTF estimation were excluded. Initial particle picking was done with template-free filament tracing with a 50-70 Å filament diameter range (Filament Tracer). Fourier-cropped particles were subjected to 2D classification, and the best classes were used to train a model for particle picking in Topaz.^59^ Particles picked by Topaz (584,291) were extracted uncropped and then curated by iterative 2D classification, ab initio reconstruction with 5 classes, and heterogeneous refinement of the 5 ab initio reconstructions, a method adapted from processing of similarly flexible, thin glycoprotein 2 filaments.^60^ Motion correction was re-performed on the best particles (99,581) with local motion correction,^61^ and an initial map (4.6 Å) was generated with non-uniform refinement.^62^ Subsequent rounds of non-uniform refinement with on-the-fly refinement of third- and fourth-order aberrations yielded a visually improved map (4.1 Å).^63^ Reference-based motion correction^64^ was performed on particles from this refinement, and a subsequent non-uniform refinement of 98,881 particles yielded the final map with 4.1 Å resolution, measured by gold standard Fourier Shell Correlation (FSC) calculated with a soft mask covering the full filament segment.

The most relevant regions of our map, e.g. the filament assembly interface, did not improve (by visual inspection) with further particle curation or processing strategies. Owing to the large helical rise, filament subunits (monomers) could be picked individually by Topaz and treated as single particles rather than filaments, and no helical symmetry was enforced at any time during data processing. The final asymmetric 3D reconstruction resolving a segment of the cN1A filament corresponding to 4 subunits was used for model building.

### Model building and refinement

The predicted structure of a cN1A monomer from the AlphaFold Database was used as an initial template for atomic modeling.^65,66^ Four copies of this template were placed into the auto-sharpened cryoSPARC map by rigid-body fitting in UCSF ChimeraX v1.12.^67^ This four-chain starting model was relaxed into the map, and the central two monomers (chains A and B) were manually fit into the map residue-by-residue with ISOLDE.^68^ The entire atomic model containing four monomers was then minimized and B-factors refined with real-space refinement in Phenix v2.0.5936.^69^ This process was repeated iteratively to improve clash score and reduce rotamer and Ramachandran outliers. Bond length and bond angle outliers identified by PDB validation were manually resolved in Coot v0.9.8.95 before model deposition.^70^ Modeling some areas of the structure was aided by simultaneous fitting into the auto-sharpened map and a map processed by EMReady2.^71^ Residues 44-212 and 249-328 of cN1A are modeled; volume for residues 1-43, 213-248, and 329-368 was not reliable enough for modeling. The final 42.4 Å helical rise and 157.3° twist values were calculated from the atomic model in UCSF ChimeraX v1.12. Buried surface area, interfacing residues, and interfacing bonds of the filament assembly interface were calculated with PISA.^36^ Directional FSC was calculated with the remote 3DFSC processing server (https://3dfsc.salk.edu/).^72^ All cryo-EM figures were prepared with UCSF ChimeraX v1.12. Data collection parameters and refinement statistics are summarized in Supp. Table 1.

## Acknowledgements

Edward Chan is gratefully acknowledged for the gift of the It2006 serum. We thank Baziel van Engelen, Benno Küsters, and Christiaan Saris for helpful discussions and advice, and Thomas Oosterhof for histochemical analyses. We thank the Arnold and Mabel Beckman Cryo-EM Center at the University of Washington for the use of electron microscopes.

## Statements and Declarations

### Author contribution

Fleur Brinkman: Investigation, Visualization, Writing – Original Draft, Methodology. John Calise: Investigation, Visualization, Writing – Original Draft, Formal analysis, Methodology. Amrah Weijn: Investigation. Josephine Molfino: Investigation. Joel D. Quispe: Investigation. Anke Rietveld: Investigation. Megan Herbert: Investigation. Annemarie van der Heijden: Investigation. Evan Spruijt: Conceptualization, Supervision, Writing – Review C Editing. Justin M. Kollman: Supervision, Funding acquisition, Writing – Review C Editing. Ger Pruijn: Conceptualization, Supervision, Writing – Review C Editing.

### Data availability

Coordinates for the atomic model of cN1A are deposited in the PDB with accession code 40KB. Cryo-EM maps are deposited in the EMDB with accession code EMD-90176. All other data are available upon request.

## Figure Legends – Supplementary Figures

**Supplementary Figure 1.**
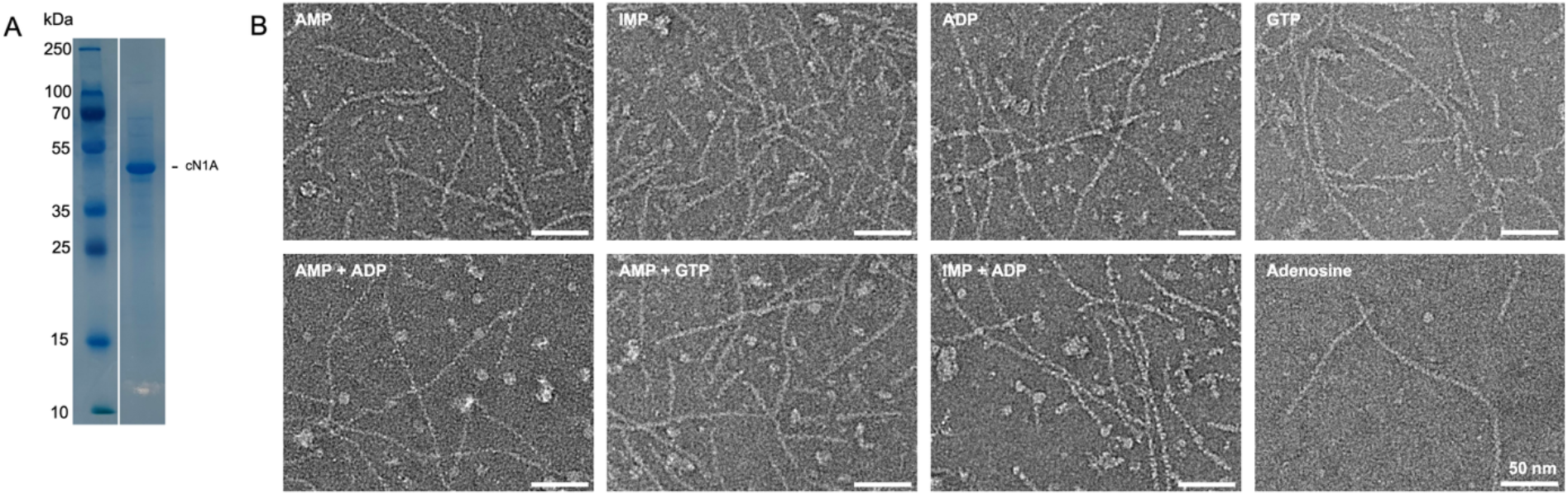
Negative stain EM of cN1A filaments in the presence of ligands. **A:** Purified recombinant His_6_-tagged cN1A protein used for *in vitro* experiments was analyzed by SDS-PAGE. Protein was stained with Colloidal Coomassie dye. The position of molecular weight markers is indicated on the left (in kDa). **B:** 2 µM cN1A was incubated with 10 mM ligand(s) (indicated in each panel) and 20 mM MgCl_2_ for 15 min at RT prior to preparation of negative stain grids. Scale bars: 50 nm.

**Supplementary Figure 2.**
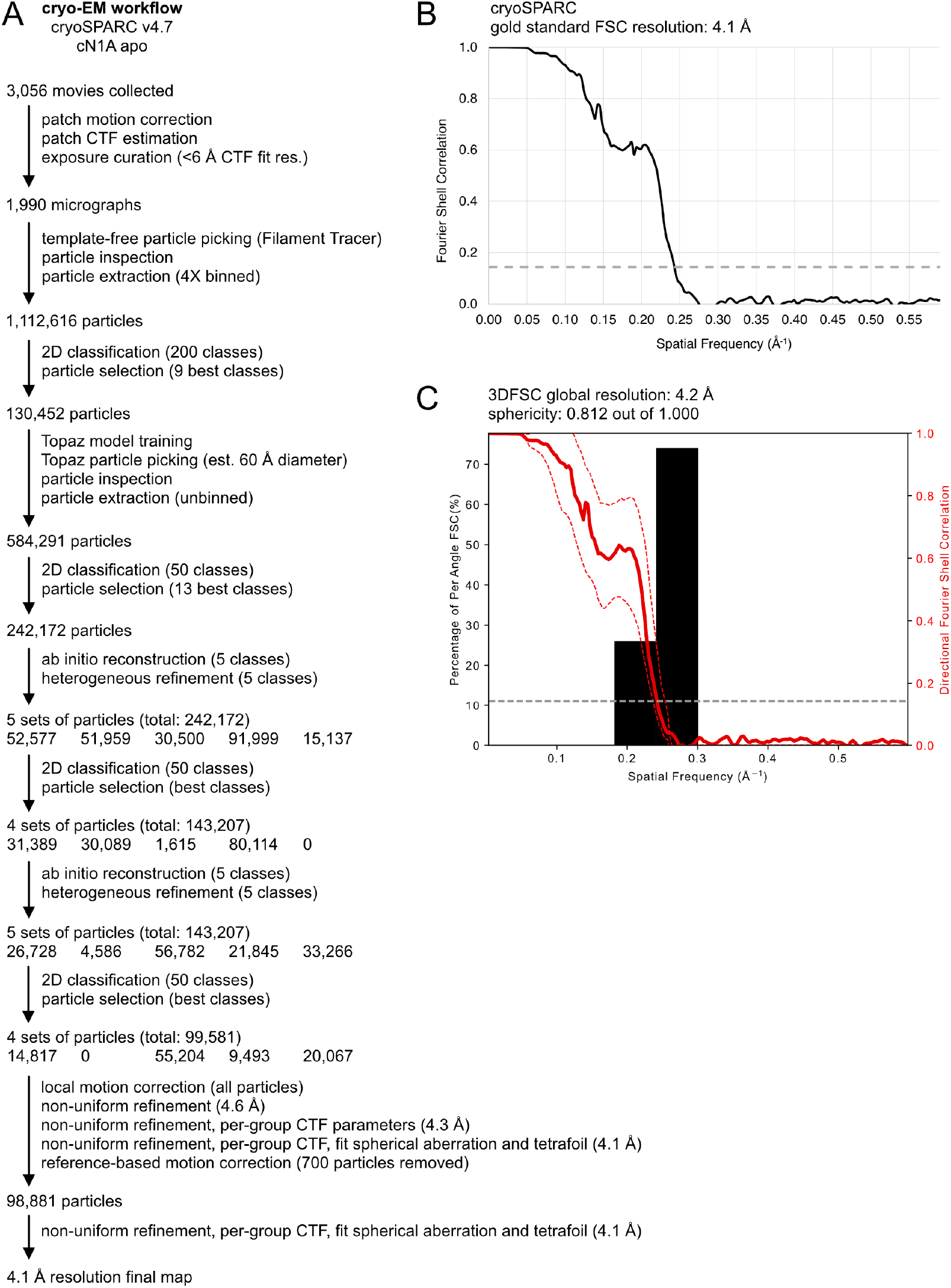
Cryo-EM data processing workflow for the cN1A filament structure. **A:** Summarized data processing workflow from cryoSPARC that resulted in the final map. Additional processing strategies and analyses that did not improve the map are not shown. **B:** Fourier shell correlation (FSC) curve for the final map, calculated in cryoSPARC using a soft mask covering the full filament segment. **C:** Directional FSC plot calculated with 3DFSC. Red dashed lines indicate ±1 standard deviation from the mean of the directional FSC.

**Supplementary Figure 3:**
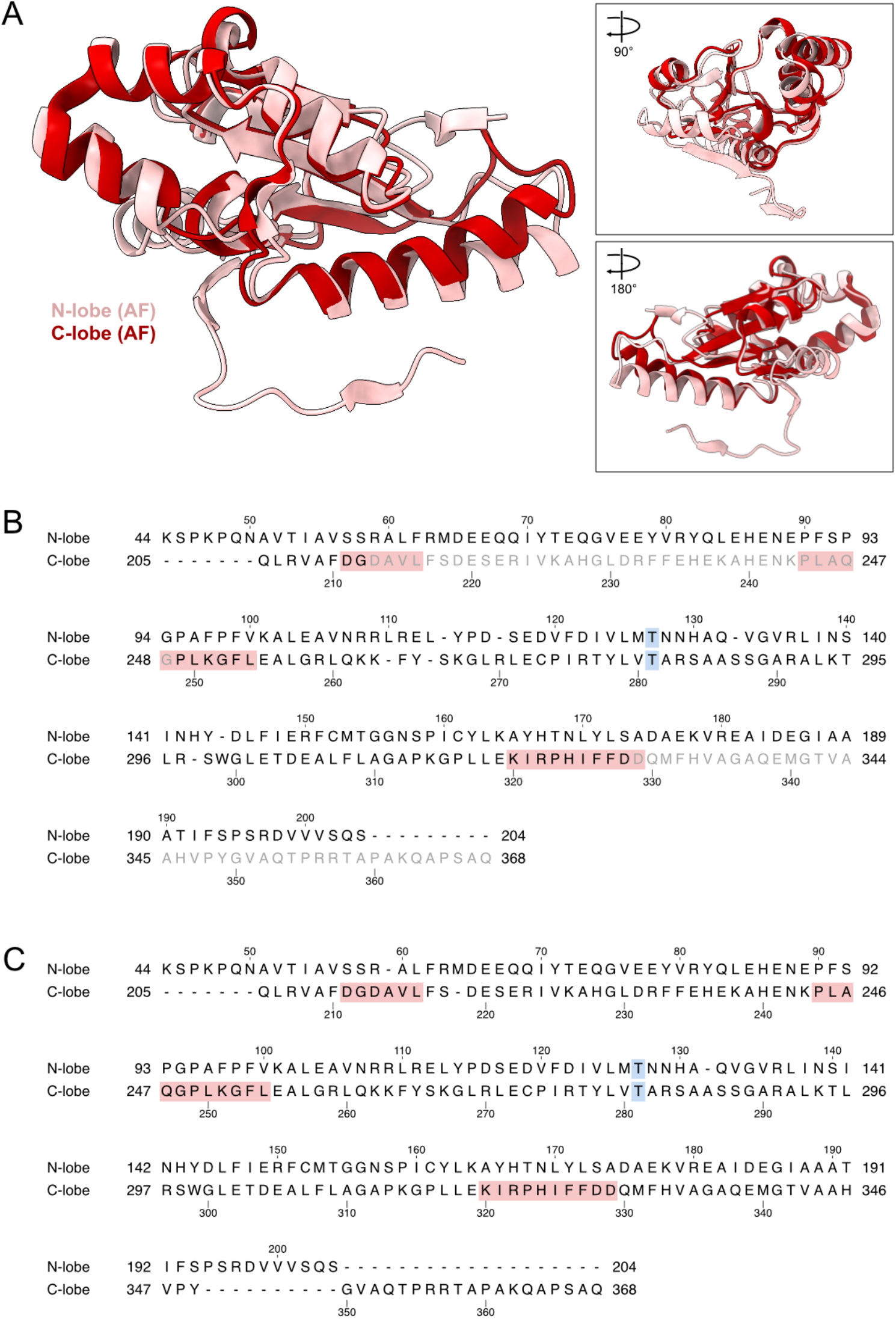
Superposition of the AlphaFold N-lobe and C-lobe and structure-based sequence alignments of the lobes. **A:** Superposition of the N-lobe (pink) and C-lobe (red) from the AlphaFold predicted structure, calculated with TM-align. **B:** Structure-based sequence alignment of the N-lobe and C-lobe calculated from our atomic model (residues 44-212 and 249-328). Residues not modeled in our structure are shown in grey for reference. These residues were not included in the structural alignment, and their positions relative to the N-lobe are shown for display only. **C:** Structure-based sequence alignment of the N-lobe and C-lobe calculated from the full-length AlphaFold predicted structure, which aligns regions that are unmodeled in our experimental structure (C-lobe 213–248 with N-lobe 59–94). Residues 1-43 have no structural counterpart and are not shown. Residue numbers are shown at the start and end of each row and at every tenth residue. Dashes indicate alignment gaps. Catalytic motif I (211-216), motif S (244-254), and motif III (320-329) are shaded red. Pseudosymmetric equivalent pair T127 and T281 are shaded blue.

**Supplementary Figure 4:**
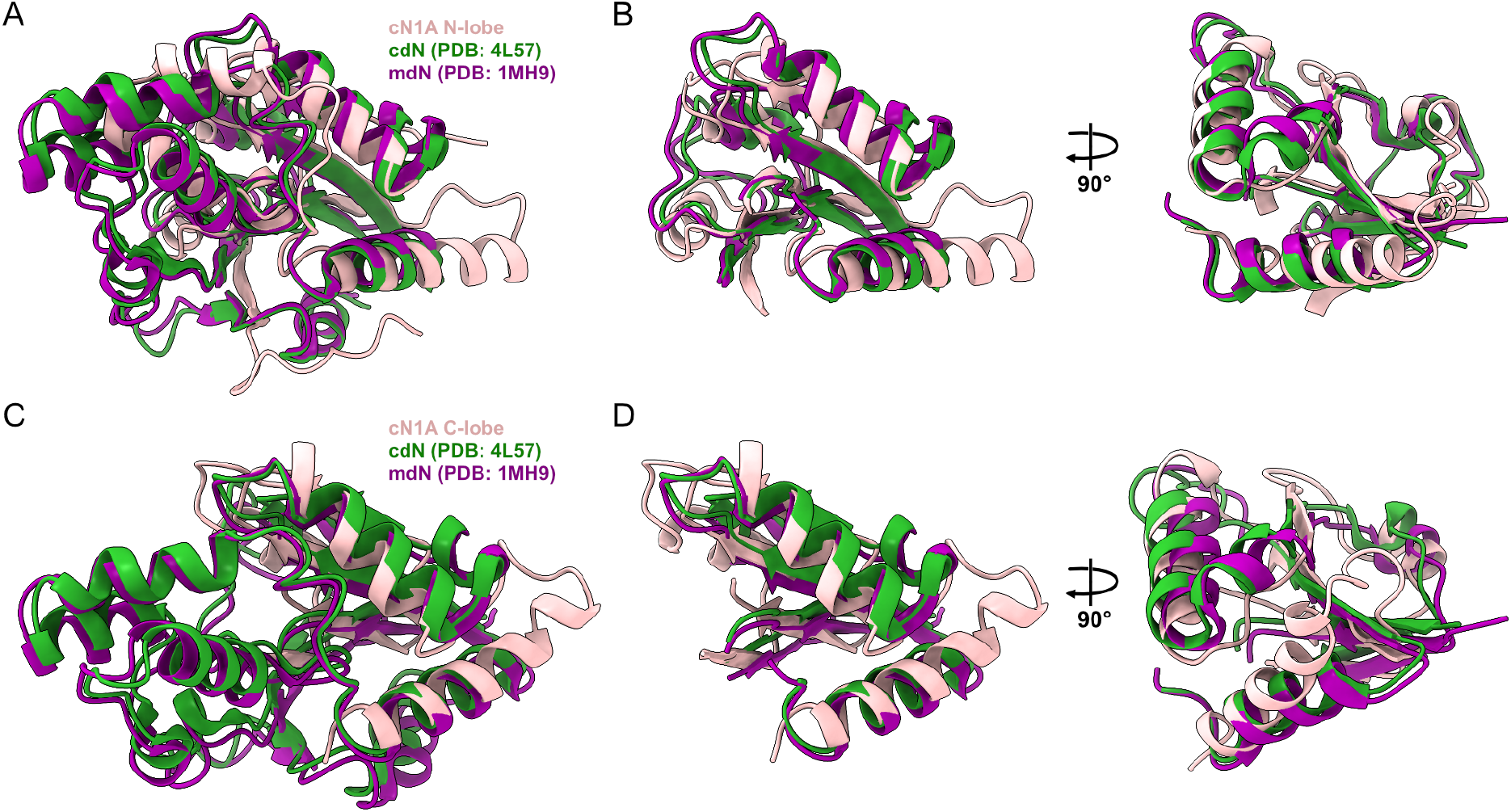
Comparison of the cN1A fold to other human nucleotidases. **A:** Superposition of the cN1A N-lobe (pink) with cytosolic 5’(3’)-deoxyribonucleotidase cdN (green; PDB: 4L57) and mitochondrial 5’(3’)-deoxyribonucleotidase mdN (purple; PDB: 1MH9). Pairwise independent structure-based alignments of cN1A with cdN and cN1A with mdN were performed with TM-align and displayed as an overlay aligned on cN1A. N-lobe vs. cdN: TM-score = 0.58, RMSD = 3.7 Å. N-lobe vs. mdN: TM-score = 0.59, RMSD = 3.7 Å. **B:** The same superposition as panel A is shown, but with the small domain and other non-aligning residues of each protein hidden to improve visibility of the highly similar core fold. Residues hidden are cN1A: 44-50,60-92,193-204; cdN: 13-75,164-195; mdN: 44-105,195-227. **C:** Superposition of the cN1A C-lobe (pink) with the same cdN and mdN models, calculated and aligned in the same manner. C-lobe vs. cdN: TM-score = 0.52, RMSD = 3.7 Å; C-lobe vs. mdN: TM-score = 0.52, RMSD = 3.2 Å. **D:** The same superposition as panel C, but with hidden residues (cdN: 13-75,145-195; mdN: 44-105,176-227).

---

**Supplementary Table 1.**
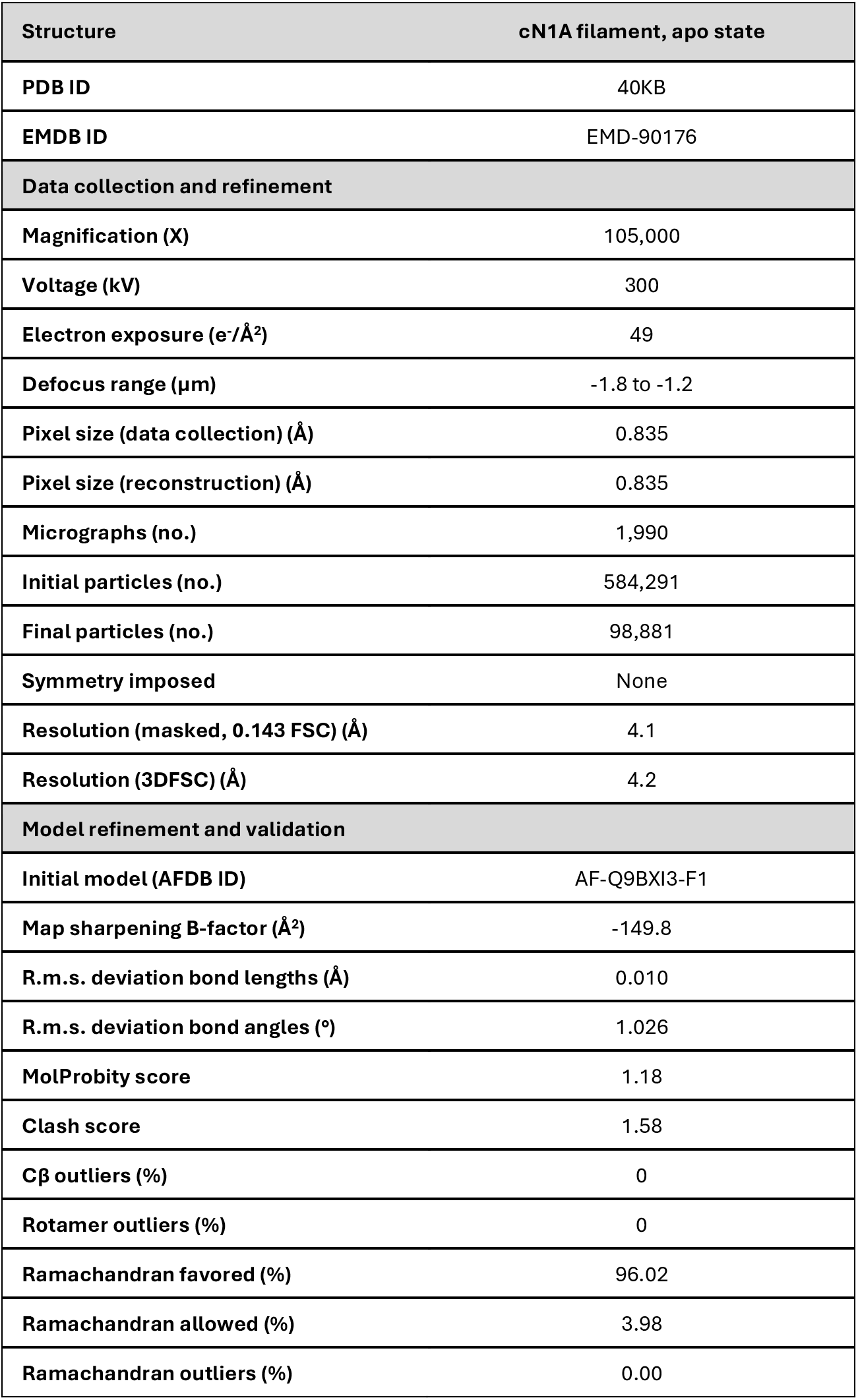
Cryo-EM data collection, refinement, and validation statistics.

| **Structure** | **cN1A filament, apo state** |
|---|---|
| **PDB ID** | 40KB |
| **EMDB ID** | EMD-90176 |
| **Data collection and refinement** |  |
| **Magnification (X)** | 105,000 |
| **Voltage (kV)** | 300 |
| **Electron exposure (e<sup>-</sup>/Å<sup>2</sup>)** | 49 |
| **Defocus range (µm)** | -1.8 to -1.2 |
| **Pixel size (data collection) (Å)** | 0.835 |
| **Pixel size (reconstruction) (Å)** | 0.835 |
| **Micrographs (no.)** | 1,990 |
| **Initial particles (no.)** | 584,291 |
| **Final particles (no.)** | 98,881 |
| **Symmetry imposed** | None |
| **Resolution (masked, 0.143 FSC) (Å)** | 4.1 |
| **Resolution (3DFSC) (Å)** | 4.2 |
| **Model refinement and validation** |  |
| **Initial model (AFDB ID)** | AF-Q9BXI3-F1 |
| **Map sharpening B-factor (Å<sup>2</sup>)** | -149.8 |
| **R.m.s. deviation bond lengths (Å)** | 0.010 |
| **R.m.s. deviation bond angles (°)** | 1.026 |
| **MolProbity score** | 1.18 |
| **Clash score** | 1.58 |
| **Cβ outliers (%)** | 0 |
| **Rotamer outliers (%)** | 0 |
| **Ramachandran favored (%)** | 96.02 |
| **Ramachandran allowed (%)** | 3.98 |
| **Ramachandran outliers (%)** | 0.00 |

